# Parallel gene expression changes in ventral midbrain dopamine and GABA neurons during normal aging

**DOI:** 10.1101/2025.02.28.640868

**Authors:** Ana Luiza Drumond-Bock, Harris E. Blankenship, Kevin D. Pham, Kelsey A. Carter, Willard M. Freeman, Michael J. Beckstead

## Abstract

The consequences of aging can vary dramatically between different brain regions and cell types. In the ventral midbrain, dopaminergic neurons develop physiological deficits with normal aging that likely convey susceptibility to neurodegeneration. While nearby GABAergic neurons are thought to be more resilient, decreased GABA signaling in other areas nonetheless correlates with age-related cognitive decline and the development of degenerative diseases. Here, we used two novel cell type-specific Translating Ribosome Affinity Purification models to elucidate the impact of healthy brain aging on the molecular profiles of dopamine and GABA neurons in the ventral midbrain. By analyzing differential gene expression from young (6-10 month) and old (>21 month) mice, we detected commonalities in the aging process in both neuronal types, including increased inflammatory responses and upregulation of pro-survival pathways. Both cell types also showed downregulation of genes involved in synaptic connectivity and plasticity. Genes involved in serotonergic signaling were upregulated with age only in GABA neurons and not dopamine-releasing cells. In contrast, dopaminergic neurons showed alterations in genes connected with mitochondrial function and calcium signaling, which were markedly downregulated in male mice. Sex differences were detected in both neuron types, but in general were more prominent in dopamine neurons. Multiple sex effects correlated with the differential prevalence for neurodegenerative diseases such as Parkinson’s and Alzheimer’s seen in humans. In summary, these results provide insight into the connection between non-pathological aging and susceptibility to neurodegenerative diseases involving the ventral midbrain, and identify molecular phenotypes that could underlie homeostatic maintenance during normal aging.

## Introduction

Biological aging is fundamentally variable. Humans and laboratory rodents exhibit great variance in cognitive abilities as they age [1–3], and individual tissues and cell types exhibit a wide range of responses to aging and its hallmarks [4–8]. Brain aging is a leading risk factor for neurodegeneration [8–10] and likely plays a causative role in the development of diseases such as Alzheimer’s (AD) [11] and Parkinson’s (PD) [12, 13]. Although findings in transcriptomic changes across the lifespan reveal selective sensitivity of regional cell populations to specific neurodegenerative diseases [7, 14, 15], detailed mechanisms behind aging-related susceptibility to neurodegeneration are not well understood, especially within vulnerable brain regions.

Neurons of the ventral midbrain that synthesize and release dopamine may be particularly susceptible to aging and have been described as “biomarkers of aging” whose functional decrements may reflect a “core mechanism of aging itself” [16]. While multiple subtypes have been described, dopamine cell bodies can be roughly divided between the substantia nigra pars compacta (SNc), which projects through the nigrostriatal pathway and is necessary for the initiation of voluntary movement, and the ventral tegmental area (VTA), which projects widely through the mesocorticolimbic pathway to regulate cognitive, motivational, and affective behaviors [17, 18]. Nigrostriatal function naturally decreases with age [19–21], and motor impairment due to dopamine neurodegeneration is a hallmark of PD [22–24]. Conversely, recent work has selectively implicated VTA dopamine neurons in the development of AD [25–27].

While subsets of dopamine neurons also noncanonically release the inhibitory neurotransmitter GABA [28], a non-dopaminergic midbrain population of vesicular GABA transporter (VGAT)-expressing neurons serves as both local interneurons and projection cells [29–33]. Although GABA neurons demonstrate relative resiliency [34], recent longitudinal studies point to a global decrease of GABA with aging in multiple regions of the brain [35]. GABA has broad effects due to its role as the brain’s principle fast inhibitory neurotransmitter, and decreased GABA signaling has been associated with the development of neurodegenerative diseases [36–38]. Particularly in AD, alterations of the balance between excitatory and inhibitory signaling could contribute substantially to cognitive decline [39]. Furthermore, a modulatory increase of GABA availability can ameliorate the effects of age in midbrain auditory neurons [40]. The intricate ways in which GABA and dopamine neurons interact in the ventral midbrain and the importance of dysfunctional dopamine release in age-linked neurodegenerative diseases makes the understanding of age effects in both neuronal populations essential.

To elucidate the impacts of aging on midbrain dopamine and GABA neurons, here we used a genetic, cell type-specific Translating Ribosome Affinity Purification (TRAP) approach to delineate the gene expression changes across the lifespan in mice and assess cellular alterations in healthy brain aging. Although we observed vast differences in the regulation of individual genes between the two cell types, the predicted age-driven biological pathways were very similar. Effects common to both neurons included upregulation of inflammatory responses and pro-survival signaling genes, and downregulation of cell connectivity related genes, associated with synaptic transmission and plasticity. These results confirm previous findings connecting aging with the susceptibility to neurodegenerative diseases [41–45], while also highlighting potential protective mechanism used by dopamine and GABA neurons to maintain homeostasis and proper brain function. Results also indicated sex-specific molecular phenotypes that could reflect the incidence of individual neurodegenerative diseases [12, 46–48]. Overall, the data provide a detailed description of age-driven molecular effects in both dopamine and GABA neurons from a single brain area (the ventral midbrain) and could be useful in targeting the biological consequences of aging in these cell populations.

## Material and methods

### Animals

Adult male and female DAT^IRES*cre*^ (RRID:IMSR_JAX:006660; [49]), Vgat-ires-cre knock-in (C57BL/6J) (RRID:IMSR_JAX:028862; [50]) and NuTRAP (RRID:IMSR_JAX:029899; [51]) mice were originally obtained from The Jackson Laboratory. These mice were group housed and mated at the OMRF Comparative Medicine facilities, on a 12h light/dark cycle. To generate “DAT;NuTRAP” mice, DAT^IRES*cre*^ (cre/cre) males were mated with homozygous NuTRAP^Flox/Flox^ females, and “VGAT;NuTRAP” mice were generated by mating Vgat-ires-cre (cre/cre) males with homozygous NuTRAP^Flox/Flox^ females (Figure 1A,B). The first two progenies of each breeding pairs were genotyped, in accordance with Jackson suggested PCR protocol, for both respective-*Cre* and *Flox* genes (see Table S1 for specific primer sequences). In addition, sporadic genotyping throughout the fertile life of each pair was performed to ensure accurate genotypes. Experimental mice were naturally aged for at least 21 months, and their transcriptomic profiles were compared to those of young adults (6-10 months old). All animal procedures were performed in accordance with the OMRF Institutional Animal Care & Use Committee (IACUC).

**Figure 1 –.**
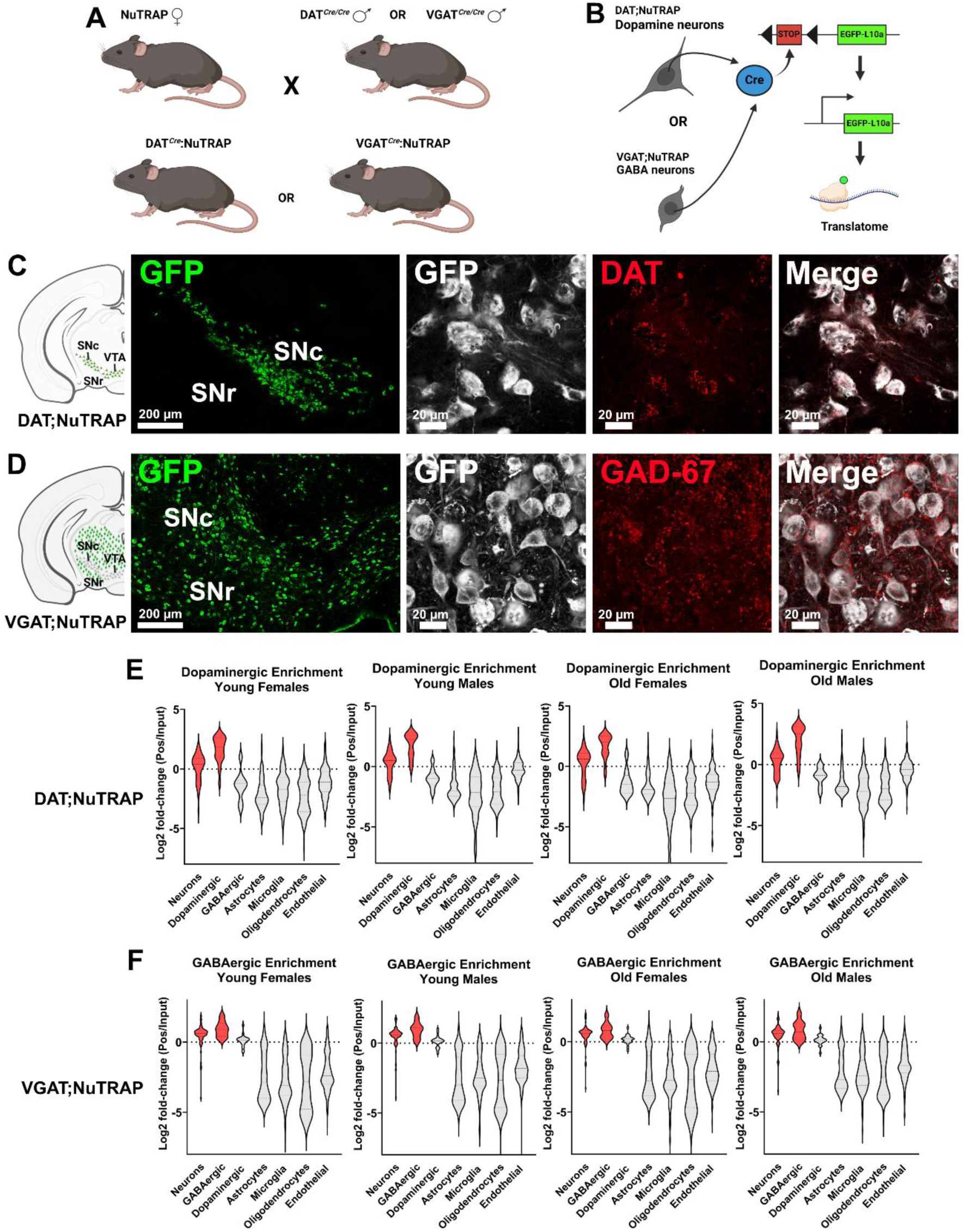
Cell type-specific NuTRAP mouse models. (A) DAT;NuTRAP and VGAT;NuTRAP mice were generated by crossing NuTRAP mice with DAT-Cre or VGAT-Cre mice. (B) In the presence of *Cre-recombinase*, the floxed stop codon was removed and ribosomal subunit L10a fused with GFP was expressed in a cell type-specific manner. (C) In DAT;NuTRAP mice, GFP signal was detected in dopamine transporter (DAT) expressing cells. (D) In VGAT;NuTRAP mice, GFP signal was detected in GAD-67 expressing cells. SNc: substantia nigra pars compacta; VTA: ventral tegmental area; SNr: substantia nigra pars reticulata. (E) Dopaminergic enrichment of the RNA-Seq dataset for DAT;NuTRAP mice, separated by age and sex. (F) GABAergic enrichment of the RNA-Seq dataset for VGAT;NuTRAP mice, separated by age and sex.

### Histology processing and immunofluorescence

Cell type specificity of the GFP-tagged ribosomes was assessed by immunofluorescence imaging (Figure 1C,D). At euthanasia, mice received an intraperitoneal injection of 2,2,2-tribromoethanol (Sigma-Aldrich, cat# T48402-25G, 0.25 g/kg). Mice then underwent cardiac perfusion with 10% sucrose (Sigma-Aldrich, cat# S7903-5KG), followed by 4% paraformaldehyde (Sigma-Aldrich, cat# 158127-500G) in 1x phosphate-saline buffer (PBS – Fisher Bioreagents, cat# BP399-1), using a Perfusion Two automated pressure perfusion system (Leica). After harvest, the entire brain was placed in 4% paraformaldehyde for 24 h, and then in 30% sucrose, and stored in 4°C until embedding. Tissues were embedded in OCT compound (Sakura, cat# 4583) and 50 µm coronal slices were sectioned with a CryoStar™ NX70 Cryostat (Thermo Scientific™). Brain slices were maintained in Cryoprotectant 3 (30% sucrose, 30% ethylene glycol, 1% PVP-40, in 0.1 M PBS) until time of staining. At staining, free-floating brain slices were washed with 1x PBS, for removal of the cryoprotectant, and then permeabilized with 0.2% Triton-X (Sigma-Aldrich, cat# T8787) in 1xPBS (PBS-T) for 1 h (4x 15 min). After PBS-T, slices were blocked for an hour with 7% normal donkey serum (NDS – Jackson Immuno Research, cat# 017-000-121) prepared in 0.2% PBS-T. Table S1 lists all antibody information and dilutions. Incubation with primary antibodies and 7% NDS was performed at 4°C for 72 h, after which samples were washed with 0.2% PBS-T for 1 h (4x 15 min). Incubation with secondary antibodies and 7% NDS was performed at room temperature (24°C) for 2 h, after which slices were washed in 0.2% PBS-T (2x 15 min) and 1x PBS (2x 15 min). Brains slices were then transferred to coated slides (Globe Scientific, cat# 1358P), mounted with ProLong glass anti-fade mounting media (Invitrogen, cat# P36980) and covered with glass coverslips (Fisherbrand, cat# 12-545). Immunofluorescence images were obtained at the Imaging Core Facility at OMRF, using a LSM 710 Confocal microscope (Zeiss). Individual channel images were consistently (for all groups) adjusted for brightness and contrast when necessary.

### Sample collection for RNA

Mice were anesthetized with isoflurane and rapidly decapitated. To minimize activity-induced transcriptomic changes, the brain was immediately removed and placed in ice-cold choline chloride cutting solution, containing 110 mM choline chloride; 2.5 mM KCl; 1.25 mM Na2PO4; 0.5 mM CaCl2; 10 mM MgSO4; 25 mM glucose; 11.6 mM Na-ascorbate; 3.1 mM Na-pyruvate, 26 mM NaHCO3; 12 mM N-acetyl-L-cysteine; and 2 mM kynurenic acid. Using a vibrating microtome (Leica VT1200s), 600 µm-thick horizontal brain slices were collected, accounting for the entire rostral-caudal length of the ventral midbrain. Sections were moved onto a pre-chilled collection block, where the surrounding tissue (such as the pons, hippocampus, and cortex) was removed. Remaining bilateral midbrain portions (containing the substantia nigra pars compacta [SNc] and the ventral tegmental area [VTA]) were collected into Eppendorf tubes and flash-frozen for RNA preservation. All samples were stored in −80°C freezer until the time of TRAP and mRNA isolation [52].

### TRAP and mRNA extraction

TRAP isolation was carried out as previously reported [52–54]. Briefly, midbrain slices were placed in 200 µl of ice-cold homogenization buffer (50mM Tris, pH 7.4; 12mM MgCl2; 100mM KCl; 1% NP-40; 1 mg/ml sodium heparin; 1mM DTT) supplemented with 100 μg/mL cycloheximide (Millipore, cat# C4859–1ML), 200 units/ml RNaseOUT™ Recombinant Ribonuclease Inhibitor (ThermoFisher, cat# 10777019), and 1× cOmplete™, EDTA-free Protease Inhibitor Cocktail (Milipore, cat# 11836170001). Homogenization used a cordless motor pestle (Kimble #749520-0090 and #749540). After initial homogenization, an additional 500 µl of buffer was added to the samples, washing the pestle in between pulses. Homogenate solution volume was then brought to a total of 1.5 ml and centrifuged at 12,000 g for 10 min at 4°C. After centrifugation, 100 µl of the supernatant was removed and set aside on ice (“input fraction”). The remaining supernatant (~900 µl of “positive fraction”) was transferred into a fresh tube and incubated with 1 µl anti-GFP antibody (Abcam, cat# ab290). Both input and positive samples were placed in an end-over-end rotating mixer and incubated for 1 h at 4°C. Pre-washed Dynabeads protein G (Invitrogen, cat# 10003D; 30 µl per sample) were then added to the positive fraction and the final solution incubated overnight (~16 h) in a rotating mixer at 4°C. Positive tubes were then placed in a DynaMag-2 magnet and supernatant removed and saved as the “negative fraction”. The magnetic beads bound with GFP-labelled polyribosomes/mRNAs, were washed with high-salt buffer (50mM Tris, pH 7.5; 12mM MgCl2; 300mM KCl; 1% NP-40; 100 μg/ml cycloheximide; 2mMDTT) three times. After the final wash, Dynabeads were separated from the GFP-labelled polyribosomes/mRNAs by adding 350 µl of Buffer RLT (Qiagen) and 3.5 µl of 2-β mercaptoethanol, proceeding with incubation in a benchtop ThermoMixer (Eppendorf) for 10 min at 22°C. The eluted solution (“positive fraction”), free of magnetic beads, was placed in a fresh tube. mRNA isolation was carried out using RNeasy Mini kit (#74104, QIAGEN), following manufacturer’s suggested protocol. Isolated mRNA was then quantified with a Nanodrop spectrophotometer (ThermoFisher, model# ND-ONEC-W) and its quality measured in a 4150 Tapestation analyzer (Agilent, model# G2992AA) using HSRNA ScreenTape (Agilent, cat# 5067-5579).

### Library construction and RNA sequencing

Directional mRNA libraries were prepared using NEBNext Ultra II Kit for Illumina (NEB, cat# E7760L) in accordance to the manufacturer’s directions, and following previously established protocol [53, 54]. In summary, each sample of both positive and input fractions (ranging 3.5-35 ng of mRNA) were individually captured by poly-A RNA using NEBNext Poly(A) mRNA Magnetic Isolation Module (NEB, cat# NEBE7490L). mRNA was then eluted from oligo-dT beads, fragmented using a thermal cycler at 94°C for 15 min, and first and second strands of cDNA were individually synthesized following manufacturer’s guidance (NEB, cat# E7760L). The final double-stranded cDNA was purified with the use of SPRISelect

Beads (Beckman Coulter, cat# B23318) and eluted in 50 µl 0.1x TE buffer. Next, adaptor ligation was performed using 100-fold dilution of the NEBNext adaptor in dilution buffer (NEB, cat# E6609L), and PCR of the ligated products was conducted using the NEBNext Ultra II Q5 Master Mix (NEB, cat#7760L) and unique index primers (NEB, cat# E6609L), both provided with aforementioned kits, in accordance to manufacture’s protocol (14 cycles). Purified libraries were then quantified using HS dsDNA Qubit kit (ThermoFisher, cat# Q33230). Library integrity, verified in a 4150 Tapestation analyzer (Agilent, model# G2992AA) using HS D1000 ScreenTape (Agilent, cat# 5067-5584), revealed an average peak size of 332bp. Libraries for each sample were then pooled at 5 nM concentration and sequenced by the OMRF Clinical Genomics Center using the Illumina NovaSeq 6000 system (S4 PE150).

### RNA-Seq analysis

Using a high-performance computing (HPC) cluster, fastq files were submitted to pre-alignment quality control (QC) analysis [55], after which adaptors were removed and reads were trimmed for poly-G and Q<20 [56]. Reads were aligned to the mouse genome assembly mm39 using STAR v2.7.10b [57, 58] and quantification was performed using featureCounts [59], and post alignment QC reports generated. Low read counts (read-counts <10) were removed from the count matrix using RStudio v.4.3.2 [60, 61]; prior to differential analysis. Normalization and differential analysis was performed with DESEq2 [62] (R/Bioconductor [61]), and Benjamin-Hochberg multiple testing correction. All remaining data processing and plot generation were performed in RStudio v.4.3.2.

### Enrichment analysis

To confirm cell type-specific enrichment of the positive fractions, DESEq2 analysis was used to compare gene expression between positive (condition) vs. input (control) fractions. Using RStudio, the resulting count matrices were filtered for a pre-defined list of genes (Table S2) for detection of neuronal [63], dopaminergic, GABAergic, astrocytic, microglial, oligodendrocytic, and endothelial specific genes [63]. Samples were separated by cell-type (dopaminergic and GABAergic), sex (males and females) and age (young and old), and each group analysis performed individually. Resulting Log2 fold-change between positive and input fractions (Figure 1E,F) was plotted using GraphPad Prism v.10 [64].

### Differential expression analysis for aging

Post-alignment raw count matrices were processed prior to differential expression analysis, using RStudio, in the following manner: 1) Removal of genes with low counts; 2) Removal of astrocytic and microglial specific genes [63] (Table S2); 3) Batch correction for (a) collection date, (b) sequencing date, (c) duplication levels and (d) millions of reads. DESeq2 normalization and differential analysis allowed for comparison of old animals (condition) vs. young animals (control). Overall aging analyses for DAT;NuTRAP (n=10 young and 9 old) and VGAT;NuTRAP (n=10 young and 8 old) were performed independently. Aging analysis was also evaluated in a sex-separate manner between old females (condition) vs. young females (control) (DAT;NuTRAP n=5 young and 4 old, and VGAT;NuTRAP n=6 young and 5 old) and between old males (condition) vs. young males (control) (DAT;NuTRAP n=5 young and 5 old; VGAT;NuTRAP n=4 young and 3 old).

### Gene ontology and Ingenuity Pathway analysis

Using RStudio, the differentially expressed genes (DEGs) for each comparison (cut off Log2FC = |0.5|; pAdj <0.5) were processed and analyzed for changes in biological processes using Kyoto Encyclopedia of Genes and Genomes (KEGG) [65] and Gene Ontology (GO) [66] [67, 68] pathway analysis. Statistical significance for KEGG and GO were p values <0.05 and q values <0.2. DAT;NuTRAP mice were further analyzed for synaptic ontology pathways using the SynGO platform [69]. DESeq2 results were also uploaded into the Qiagen Ingenuity Pathway Analysis [70] for further understanding of upregulated and downregulated pathways, and for comparison between gene regulation in different groups (females vs. males and DAT;NuTRAP vs VGAT;NuTRAP).

## Results

### NuTRAP crosses successfully provide translatomes enriched for midbrain cell types

To verify that DAT;NuTRAP and VGAT;NuTRAP mice express GFP ribo-tags in a cell type-specific manner, we stained PFA-fixed midbrain slices with antibodies targeting GFP and DAT, in DAT;NuTRAP mice, or GAD-67 (glutamate decarboxylase, the enzyme essential for synthesis of GABA [71]) in VGAT;NuTRAP mice. In DAT;NuTRAP mice, we observed GFP staining in areas corresponding to the SNc and VTA (Figure 1C) and as expected the GFP signal overlapped DAT puncta in dopaminergic neurons. We observed a broader staining of GFP in VGAT mice, as expected given the multiple types of GABA neurons present in the ventral midbrain (Figure 1D). The GFP signal in VGAT;NuTRAP mice also overlapped with the GAD-67 staining present in the cell bodies and processes of GABA neurons.

Differential analysis of the RNA-seq data between positive and input fractions for all DAT;NuTRAP groups (young females or males, and old females or males) confirmed that the translatomes isolated from these mice were successfully enriched for dopaminergic neuronal transcripts (Figure 1E and Table S2). In accordance, differential analysis performed in VGAT;NuTRAP groups also demonstrated that samples from all four groups exhibit an enrichment of neuronal and GABAergic genes (Figure 1F and Table S2). Enrichments showed a consistent pattern across ages and sexes. The immunostaining and enrichment analyses are consistent with what has been described in the literature during model validation using NuTRAP mice or dopamine neuron-specific translatome analyses [4, 51, 53]. Thus we were confident the transcriptomics datasets are enriched for dopaminergic neurons (DAT;NuTRAP) or GABAergic neurons (VGAT;NuTRAP) and proceeded with differential analysis of aging and sex-biased gene expression.

### Orthogonal correlation with public datasets

In order to validate our findings on DAT;NuTRAP and VGAT;NuTRAP neuronal aging, we compared our transcriptomics data with previously published datasets obtained both with the TRAP technique [4] and other transcriptomic techniques [14]. To validate the DAT;NuTRAP dopaminergic enrichment we used published data from Kilfeather and colleagues (https://spatialbrain.org; [4]), which used the correlation between TRAP RNA-seq and spatial transcriptomics to characterize the translatome profile of dopaminergic neurons in the midbrains of young and old mice. Spatial transcriptomics allowed for a detailed detection of gene expression changes in subpopulations of dopaminergic neurons and showed a high correlation with TRAP data. However, the TRAP approach demonstrated a greater detection power, offering the most inclusive and sensitive measure of dopaminergic gene expression. We started by analyzing our dopaminergic enrichment results (DESeq2: GFP positive fraction x bulk input RNA-seq) with the same cut off values (lfcThreshold = log_2_|1.05|), pAdj <0.01) [4]. We found an overlap of 65% among the genes with expression above the threshold (lfc > log_2_(1.05); Figure 2A; Table S3), accounting for 3,049 genes in our DAT;NuTRAP positive fraction. We determined that the majority of the missing genes (35% of Kilfeather et al. enrichment) were removed from our dataset during low-count removal (read-counts <10) and therefore reflect the exclusion criteria used in this study. Comparing just the enriched transcripts with adjusted p value < 0.01 (Pos>Input) (Table S3), we found an agreement of 97.5% of the genes showing significantly enrichment (lfc >log_2_(1.05), pAdj <0.01) in both datasets (Figure 2B). After confirming that the majority of the dopaminergic genes in our dataset agreed with those reported by Kilfeather et al, we compared the age-related changes represented in both datasets to perform independent GO and KEGG pathways analysis. Overlapping GO and KEGG terms (Figures 2C,D; Table S3) revealed agreement between changes detected in DAT;NuTRAP and some of the terms reported for protein-protein interaction of aging related DEGs [4], among them axonal extension, synaptic vesicle (endocytosis) and protein ubiquitination. Finally, we detected that all of the mitochondrial-related genes reported as downregulated [4] are also downregulated in our dataset (Tables S3, S5). Taken together, this analysis increased confidence in our dopaminergic aging dataset as it correlated closely with results from prior literature.

**Figure 2 –.**
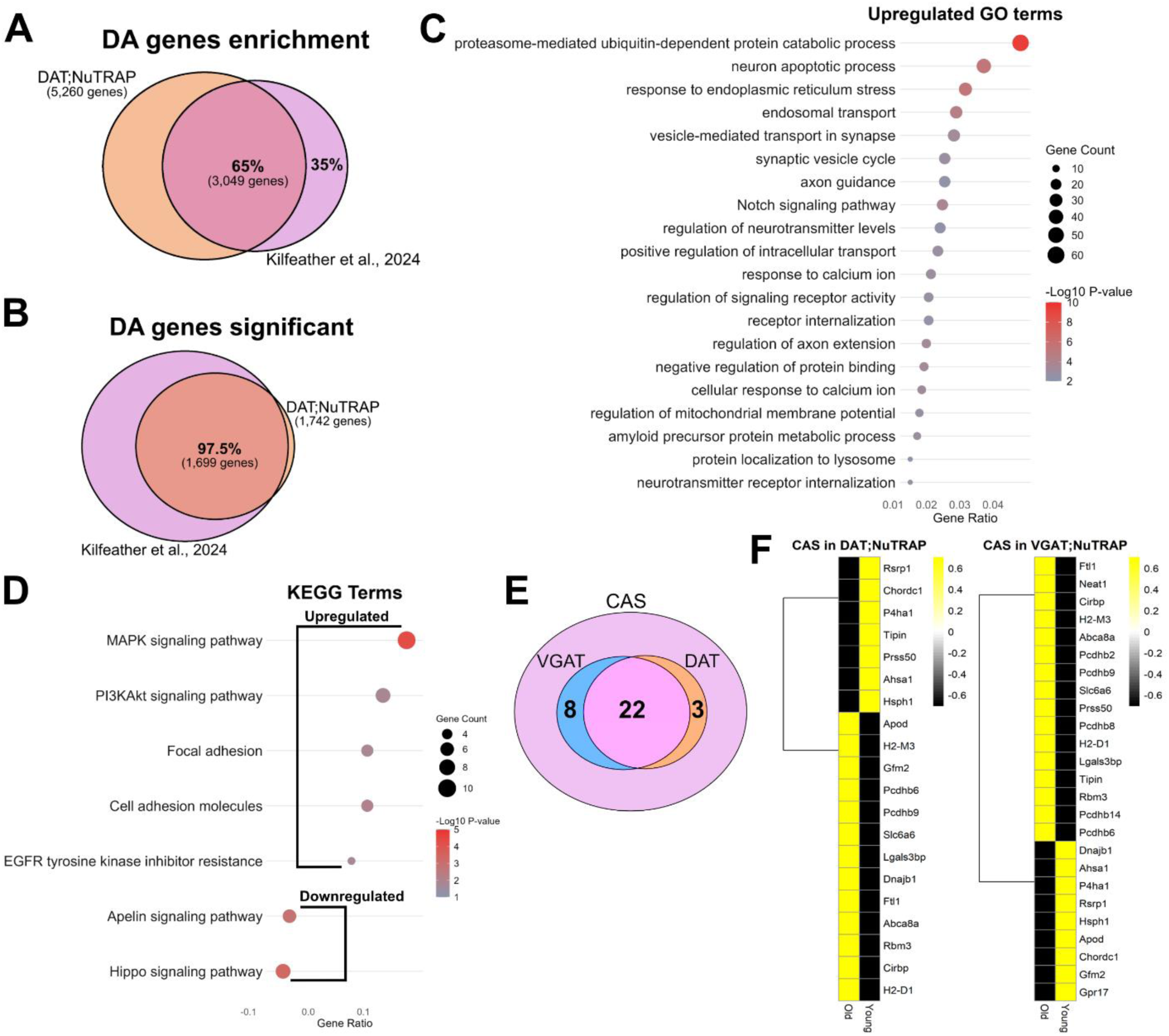
Orthogonal validation. (A) Comparison of dopaminergic gene enrichment between Kilfeather et al. (2024) and our dopamine dataset (all ages and sexes combined). (B) Overlap of significantly enriched genes between our dopaminergic dataset and Kilfeather et al. (2024). (C) Gene Ontology (GO) terms of commonly upregulated genes with age (between our data set and Kilfeather et al. (2024)). (D) Kyoto Encyclopedia of Genes and Genomes (KEGG) terms of commonly up- and downregulated genes with age (between our data set and Kilfeather et al. (2024)). Pathways analysis cut off: p value <0.05 and q value <0.02. (E) Number of “CAS” (Common Aging Score) [14] genes present in the DAT;NuTRAP and VGAT;NuTRAP datasets, and number of genes common to both groups. (F) CAS genes up- (yellow) and down- (black) regulated in DAT;NuTRAP and VGAT;NuTRAP. Cut off: Log2FC > |0.5|; Adjusted p value < 0.5.

Using a combination of region-specific bulk RNA sequencing, spatial and single-nucleus transcriptomics, Hahn and colleagues [14] created a detailed atlas of the mouse brain during aging analyzing several different brain sections. While aging-driven changes in gene expression are specific for each area of the brain, these authors detected the existence of a shared aging signature, termed “Common Aging Score - CAS”, which was present in all regions to varying degrees. Although a study of the midbrain was not included in this work, we hypothesized that a similar effect might occur in the area. We first removed microglial and astrocytic specific genes (Table S2 - 42 in total) [63] from the CAS gene set (Table S4). Next, we intersected the CAS with genes that are expected to be expressed in young dopamine and GABA neurons of the midbrain according to the Allen Brain Cell Atlas (https://portal.brain-map.org; [73]) (Figure 2E; Table S4). We found 25 CAS genes expressed in dopamine neurons and 30 in GABA neurons, with 22 of the CAS genes expressed in both cell types. From this resultant CAS gene list (40 genes) (Table S4), we detected changes in the expression (old x young) of 20 out of 25 genes in our DAT;NuTRAP dataset and 25 out of 30 genes in the VGAT;NuTRAP (Figure 2F; Table S4). There was upregulation of 13 CAS genes in DAT;NuTRAP and 16 in VGAT;NuTRAP, accounting for a large majority of the CAS expected in both cell types. Among downregulated CAS genes, 5 genes were predicted to be present in dopamine and GABA cells (Table S4; ABC: https://portal.brain-map.org; [73]) and were commonly downregulated in both DAT- and VGAT-expressing cells (Figure 2F; *Asha1*, *P4ha1*, *Rsrp1*, *Hsph1* and *Chordc1*), thus are likely to represent a common effect of aging in these cell types. Overall, our analysis detected that dopamine and GABA neurons of the midbrain are subjected to some of the signature alterations in CAS genes, in accordance with observations in other brain nuclei [14].

### Differences and similarities of the aging process in midbrain dopamine and GABA neurons

Defining DEGs with aging as an adjusted p value <0.5 and log2FoldChange > |0.5|, we detected that age promoted upregulation of 180 genes and downregulation of 212 genes in dopamine neurons (Figure 3A; Table S5) and were consistently changed across all aged samples (Figure 3B). KEGG analysis revealed that signaling pathways connected to inflammatory responses such as MAPK, Notch and PI3K/Akt were all among the most upregulated terms with aging (Figure 3C). Activation of MAPK signaling induces degeneration of dopaminergic neurons in the SNc [75] and is upregulated in AD and PD patients [76]. Notch signaling is activated in AD mouse models [77] and mutations involving Notch protein have been identified in AD patients [78]. The PI3K/Akt signaling pathway is involved in different neuronal processes. Particularly under induced stress, it exerts a cell-protective effect, enhancing expression of inflammatory cytokines and pro-survival signaling cascades [79, 80]. The activation of these pathways may suggest a neuronal response to increases in local, age-related inflammation, a phenomenon well studied in cortex and hippocampus [81–83]. Furthermore, KEGG analysis in combination with GO pathway analysis (Figure 3D) showed upregulation of genes involved in cell adhesion and transmembrane signaling, indicating enhanced responses to the extracellular environment. Finally, the dataset points toward upregulation in genes connected with glycolysis and gluconeogenesis (Figure 3C), which in combination with the aforementioned decrease in mitochondrial-related genes, suggests an alteration in dopaminergic neuron metabolic regulation [84, 85]. Aging-related mitochondrial dysfunction in the brain is a widely studied topic [85–88] and is one of the main factors connecting aging of dopaminergic neurons and the occurrence of PD [89–91].

**Figure 3 –.**
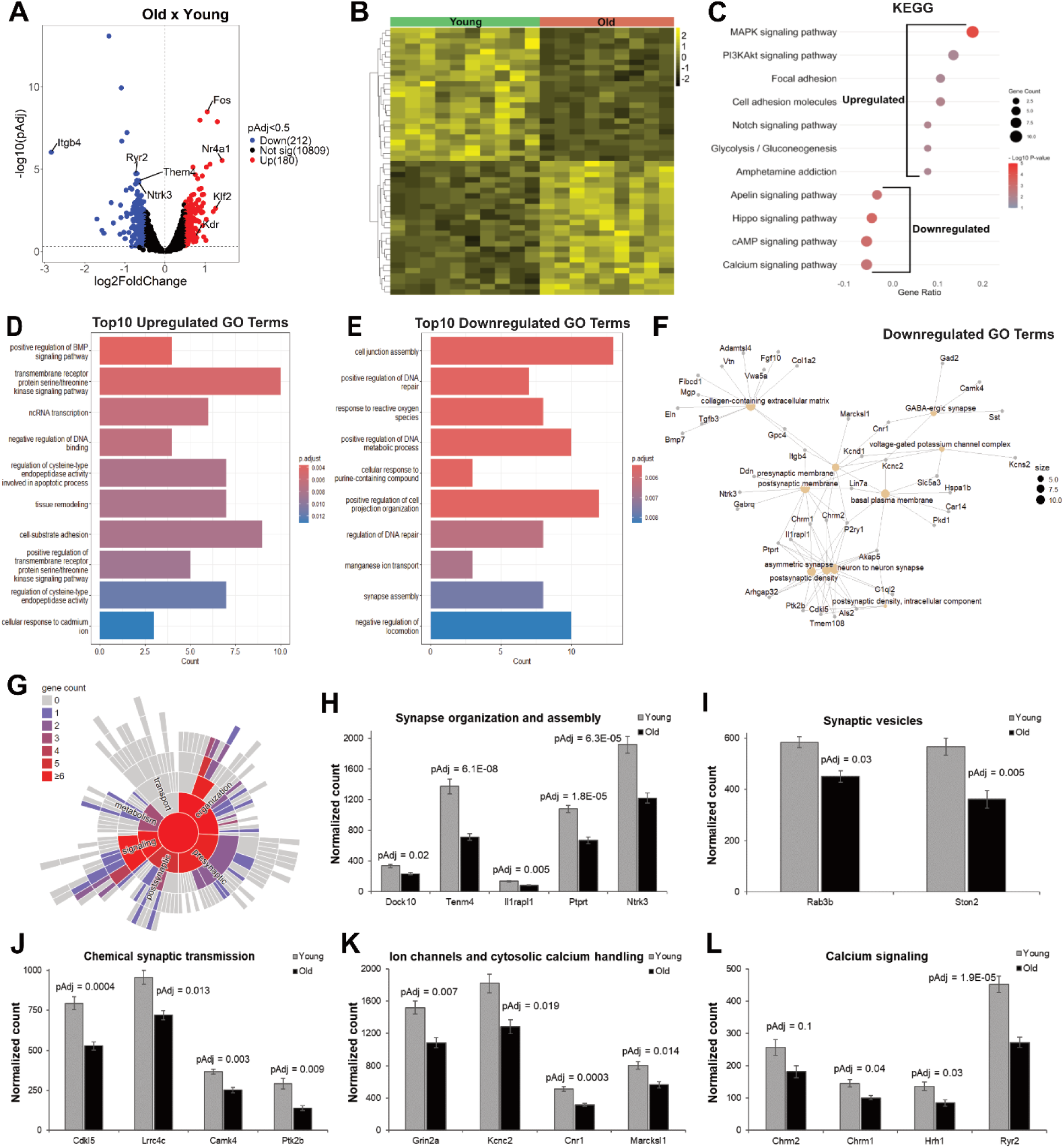
Gene expression changes with age in DAT;NuTRAP mice. (A) Volcano plot and (B) heatmap of differential expression analysis for DAT;NuTRAP mice. (C) Kyoto Encyclopedia of Genes and Genomes (KEGG) pathway analysis showing most enriched terms in DAT;NuTRAP mice. Top Gene Ontology (GO) terms of upregulated (D) and downregulated (E) genes in DAT;NuTRAP mice. (F) Gene-network analysis of downregulated genes in DAT;NuTRAP mice, showing several synaptic-related terms. (G) SynGO (Synapse Gene Ontology) analysis of synaptic-related DEGs in DAT;NuTRAP mice. (H-K) Synaptic-related genes and (L) calcium signaling-related genes significantly downregulated with age (Adjusted p value < 0.5).

Concerning downregulated terms, KEGG analysis revealed an expressive downregulation of genes involved in calcium and cAMP signaling, as well as hippo and apelin pathways (Figure 3C). Apelin plays an important role in various neurological disorders [42]. Particularly in the SNc, apelin was shown to provide a protective effect against PD in rodent models [92, 93], and potential signaling downregulation suggests an increased vulnerability of the dopaminergic neurons with aging. KEGG terms in combination with GO pathways analysis (Figure 3E) pointed to downregulation of DNA repair pathways as well as response to reactive oxygen species, providing additional insights into age-induced mechanisms of susceptibility in these neurons. A further look into the GO terms via network plot revealed downregulation of several synapse-related genes that participate in synaptic membranes, synaptic density and transmission (Figure 3F). To examine this closer, we queried a list of 23 downregulated genes using the Syngo portal (syngoportal.org, [69]; Figure 3G). We detected that the majority of synapse-related genes pointed to downregulation of terms associated with synapse organization and assembly (Figure 3H,) as well as crucial genes involved in dopaminergic transmission such as *Rab3b* [94] (Figure 3I) and *Cdkl5* [95](Figure 3J). Furthermore, several of the downregulated synaptic genes were involved in calcium regulation (Figure 3H-K). In fact, we detected seven overlapping genes between downregulated synaptic genes and the ones identified by KEGG analysis as involved in calcium signaling (Figure 3C): *Ntrk3*, *Camk4*, *Lrrc4c*, *Ptk2b*, *Grin2a*, *Kcnc2*, *Cnr1* and *Marcksl1*. In addition, we also detected downregulation of important intracellular calcium regulators such as *Chrm1* [96] and *Ryr2* [97], pointing toward a change in intracellular calcium dynamics, a neuronal age effect widely described in the literature [19, 43, 44, 98, 99]. Curiously, inhibition of ryanodine receptors (Ryr) provides a protective effect in situations of intracellular calcium dyshomeostasis [100] and could be identified as a potential self-protective mechanism against increases in free intracellular calcium, although its loss also affects synaptic plasticity by impairing remodeling of dendritic spines and decreasing excitatory synapses [97]. Overall, our analysis of age-related transcriptomic changes in dopaminergic neurons provided several insights into mechanisms of susceptibility and vulnerability of dopaminergic neurons such as the upregulation of inflammatory signaling and downregulation of mitochondrial genes and genes involved in synaptic transmission and plasticity. However, it also identified upregulation of pro-survival signaling, as well as changes in metabolic regulation and intracellular calcium dynamics, which are consistent with maintenance of homeostasis in the face of age-related deficits.

To determine if GABA neurons share similar responses to aging as dopaminergic neurons, we also probed their translatomic profile. While analyzing age-driven changes in VGAT;NuTRAP mice, we detected upregulation of 119 and downregulation of 77 genes (Figure 4A,B; Table S3). Similar to results from DAT;NuTRAP mice, KEGG analysis returned terms associated with the upregulation of pathways involved in inflammation response (Figure 4C) such as cytokine receptor interaction, TGFbeta, Notch signaling, and necroptosis. Some of the upregulated genes (Figure 4D; Table S5) were involved in serotonergic synaptic signaling, including *Tph2*, *Pla2g4e,* and *Cyp2d22*, which is upregulated with age in other areas of the brain [101]. Some of the genes were also connected to stress induction [102] and inflammation [103, 104], which could warrant further investigation. Furthermore, KEGG analysis pointed towards downregulation of genes involved in Rap1 and Ras signaling, calcium signaling, and focal adhesion (Figure 4C, E). Similar to what we observed in DAT;NuTRAP mice, these terms are associated with synaptic transmission and plasticity [105, 106]. In particular, we found that *Rasgrp1* (Figure 4E), which is downregulated with age in the human cortex [15], is also downregulated with age in dopamine neurons (Figure 2G) and is downstream from calcium signaling [105, 107], which was found downregulated in both cell types (Figures 3C and 4C).

**Figure 4 –.**
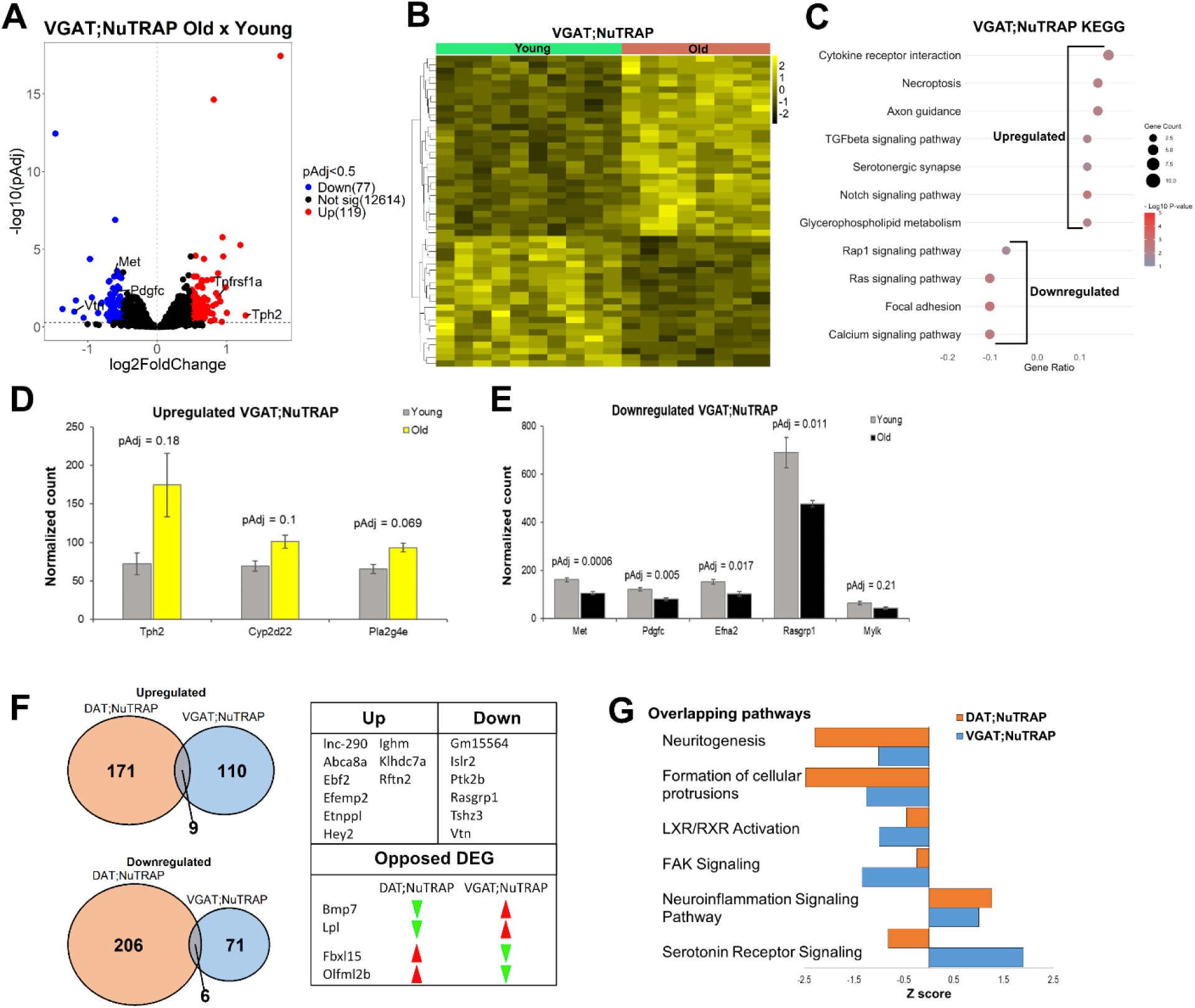
Gene expression changes with age in VGAT;NuTRAP mice and comparison between DAT and VGAT-expressing neurons. (A) Volcano plot and (B) heatmap of differential expression analysis for VGAT;NuTRAP mice. (C) Kyoto Encyclopedia of Genes and Genomes (KEGG) pathway analysis showing most enriched terms in VGAT;NuTRAP mice. Serotonergic-related genes significantly upregulated (D) (Adjusted p value < 0.5), and cell signaling-related significantly downregulated genes (E) in VGAT;NuTRAP mice. (F) Intersected up- and downregulated genes with age between both DAT;NuTRAP and VGAT;NuTRAP datasets. (G) Ingenuity Pathway Analysis (IPA) of overlapping pathways for both groups. Pathways analysis cut off: p value <0.05 and q value <0.02.

Comparatively, dopamine and GABA neurons exhibited divergent sets of individual DEGs with aging (Figures 4F), suggesting that these neurons establish unique age-related molecular phenotypes (Figure 3A and 4A; Table S5), as evidenced by the single digit number of shared up- and downregulated genes (Figure 4F). One noteworthy difference between GABA and dopamine neurons is the predicted activation of serotonin receptor signaling pathway in GABA neurons, in opposition to the predicted decrease in dopamine neurons (Figure 4G). However, the Z-scores for activated and inhibited overlapping pathways (Figure 4G) suggest that the majority of the predicted biological changes in both neuron types remain the same: activation of neuroinflammation signaling pathways and decrease in pathways that correlate with synaptic function. Taken together, we conclude that aging promotes alterations in gene expression in midbrain dopamine and GABA neurons that are very different at first look (when considering DEGs), but that converge to parallel predicted biological pathways. Overall, common translatomic changes of aged midbrain dopamine and GABA neurons reflect an increased response to inflammatory and extracellular signaling, accompanied by a decrease in synaptic transmission and plasticity. These responses correlate with neurodegeneration as well as the development of neurodegenerative diseases that have age as a risk factor [76, 78, 80, 83, 108, 109]. Furthermore, shared events such as the increased pro-survival signaling and changes in regulation of metabolism and calcium dynamics suggest that alterations in both neuronal types could be interpreted as an attempt of the cells to re-establish homeostatic levels of cellular function and neuronal connectivity during aging [44, 110].

### Sex differences in age effects on gene expression

We next sought to investigate whether some of the modifications in either cell type from DAT;NuTRAP and VGAT;NuTRAP mice were differentially influenced by sex. Differential expression analysis in DAT-females showed 681 upregulated and 588 downregulated genes (Figure 5A and Table S5) that in general were consistently changed across all aged samples (Figure 5A). In DAT-males, there were 795 genes upregulated and 834 downregulated (Figure 5B and Table S5), also consistent across samples (Figure 5B). Out of 1,400 potentially upregulated genes (males + females), only 76 genes were shared between the two groups (Figure 5C). A similar trend was observed for downregulated genes, where there was an overlap of 71 out of 1,351 genes (males + females; Figure 5C). This result points toward substantial differences in dopamine neuron aging between sexes. Of the 1,269 DEGs identified in females, only 27 were present on the X chromosome, while in males 48 of the total 1,629 DEGs were located on the X chromosome. No Y-linked DEGs were observed in either dataset. Taken together, these results suggest that the large majority of the differences in gene expression between DAT-males and DAT-females are present on autosomal genes, and are not due to sexual chromosome expression differences.

**Figure 5 –.**
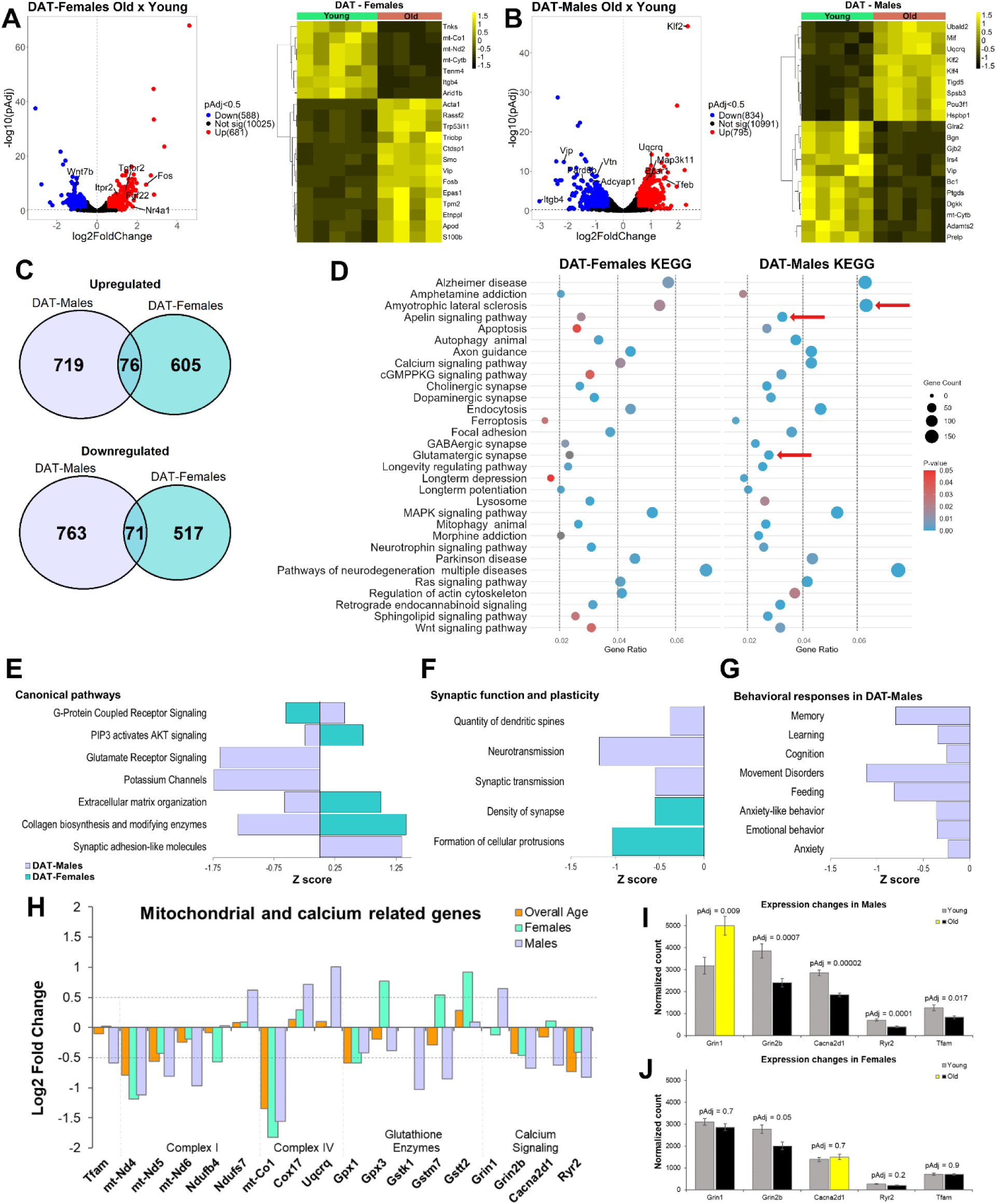
Sex-separate expression changes in dopamine neurons with age. (A) Volcano plot of differential expression analysis for females and heatmap of the top differentially expressed genes (DEG; Cut off: Log2FC > |0.5|; Adjusted p value < 0.5). (B) Volcano plot of differential expression analysis for males and heatmap of top DEGs. (C) Intersected up- and downregulated genes with age, between males and females. (D) Overall KEGG pathway analysis showing the most represented terms in both groups. The use of the same scale allows for comparison between predicted changes in both females and males. Red arrows: Different changes in pathways. (E) IPA shows comparison of activated and deactivated paths for canonical pathways of both groups. (F) IPA analysis show comparison of the downregulated pathways involved in synaptic function and plasticity for both groups. (G) IPA analysis shows predicted behavioral pathways downregulated for males, which were not altered in females. Pathways analysis cut off: p value <0.05 and q value <0.02. (H) Expression changes with age of mitochondrial-related and calcium signaling-related genes for overall aged (DAT;NuTRAP), males DAT;NuTRAP and females DAT;NuTRAP. Dotted lines show cut off values for gene expression changes (Log2FC > |0.5|). (I) Expression changes of specific genes in males, showing increased (yellow) or decreased (black) expression with age, and expression changes of the same genes in females (J). Adjusted p value (pAdj) is represented for each gene (Significant changes were defined as having pAdj < 0.5).

Despite the apparently large differences in age-driven gene expression changes for males and females, KEGG analysis performed on all DEGs in each group (Figure 5D) revealed major overlap in the biological processes affected by aging, with similar gene ratios and p values. The top affected pathway in both groups was for genes connected with neurodegenerative diseases. There were, however, noteworthy differences in the pathway analysis, particularly in males. There was a higher difference in the expression of genes associated with amyotrophic lateral sclerosis (ALS), a disease known to be connected with age [46, 111] and is more prevalent in men [112]. The apelin pathway was also significantly enriched in males (p = 5.21E-05) and was one of the most downregulated pathways in the overall DAT;NuTRAP analysis (Figure 3C). As mentioned above, apelin signaling is affected by aging [113], is connected with several neurological disorders [42], and is neuroprotective in mouse models of PD [92, 93]. Finally, genes associated with glutamatergic synapses were apparently more altered in males than in females (Figure 5D), an age-related sex effect that has been previously reported [114].

Ingenuity Pathway Analysis (IPA) of predicted activated or deactivated pathways (p <0.05, pAdj <0.5) revealed at least four canonical pathways with opposite effects between males and females (Figure 5E). The analysis predicted a male-biased downregulation of glutamate receptors (as anticipated by KEGG analysis, Figure 5D) and potassium channels. Z-score results also revealed that synaptic function and plasticity appear to be downregulated in both males and females (Figure 5F), but the specific mechanisms involved may be different between the two groups. While in males there was a predicted downregulation of quantity of dendritic spines and neuronal/synaptic transmission, changes in females were associated with decreased synaptic density. Lastly, aged males demonstrated changes in gene expression connected with the downregulation of several behavioral responses (Figure 5G), most of which have been associated with dopamine and may decline in aging individuals [21, 115]. This includes behavioral responses associated with movement disorders, calling to mind the hallmark symptoms of PD [22, 116].

One final difference of note between male and female DAT;NuTRAP mice involves mitochondrial dysfunction and calcium signaling (Figure 5H-J). While all three datasets (overall aged, aged males and aged females) showed mitochondrial dysfunction and alterations in calcium signaling as an effect of age, the changes were most prominent in males. Specifically, males showed a significant decreased in the expression of *Tfam*, the protein responsible for stabilizing and transcribing mitochondrial DNA [117], and whose knockout in dopamine neurons drives parkinsonian phenotype due to alterations in mitochondrial respiratory chain [118, 119]. Downstream of *Tfam* regulation, we observed that several mitochondrial genes involved in complex I (Figure 5H) were downregulated, and although the changes were observed in all three datasets, it was more accentuated in males. Deficiencies in mitochondrial complex I in dopamine neurons of the SNc are commonly seen in PD patients and animal models [90, 120, 121]. Interestingly, we also observed in males an increased expression of genes involved in complex IV (Figure 5H), which could be interpreted as an attempt to provide a compensatory effect to the deficiency in complex I. This observation has been seen in other studies that detected that male mice are able to maintain an overall homeostasis of synaptic mitochondrial function, despite alterations in mitochondrial proteins and bioenergetics [88]. Furthermore, males displayed downregulation of several genes involved in glutathione metabolism while females surprisingly showed upregulation of the same genes (Figure 5H). Glutathione is an intracellular antioxidant, with important protective function in dopaminergic neurons [122], and its decrease is also observed in PD patients [123]. These differences in glutathione metabolism genes suggest that female dopamine neurons likely have better mechanisms to handle oxidative stress than in males. In combination, these results suggest that dopamine neurons of aged males are more susceptible to mitochondrial dysfunction than in aged females, due to differences in several genes. Because mitochondria are critical to the homeostatic regulation of Ca^2+^ in dopamine neurons [124, 125], we looked further into the effects of age and observed more alterations in genes involved in calcium signaling in males than in females (Figures 5H-J). This correlates with previously reported observations that age-driven alteration of calcium dynamics are more relevant in males [19, 20], and altogether these data suggest that age-related calcium dynamics are connected to mitochondrial dysfunction and decreased ability of calcium handling in dopaminergic neurons in aged males.

Overall, these results identify sexual divergences in dopamine neuron gene expression with age that correlate with the prevalence of certain neurodegenerative diseases, particularly PD. Given the potential link to neurodegenerative processes, this information could be used to drive specific hypotheses involving the role of whole brain aging and its involvement in age-related neurodegeneration. Furthermore, sex biases should be taken into consideration while studying any age effect on dopaminergic circuits, behavioral processes, and related diseases.

### Sex differences in gene expression changes in GABA neurons

The last step in our analysis was to investigate whether the differences observed in aged dopamine neurons were also evident in GABA neurons (i.e., in VGAT;NuTRAP mice). Differential expression analysis in old females revealed upregulation of 252 and downregulation of 221 genes (Table S3), and that the top DEGs were consistently changed across samples (Figure 6A). In males, we observed 451 upregulated and 347 downregulated genes (Table S5), with the top DEGs again changing consistently across samples (Figure 6B). Of these, females and males shared just 36 common upregulated genes (Figure 6C) and 29 common downregulated genes, again suggesting prominent sex effects at the level of individual DEGs. Out of all age-related DEGs in VGAT-males, 33 were X-linked genes, while in VGAT-females 6 genes were X-linked genes, with an overlap of 4 common genes between both groups. This suggests that the majority of sex effects in gene expression were autosomal and not associated with sex chromosomes.

**Figure 6 –.**
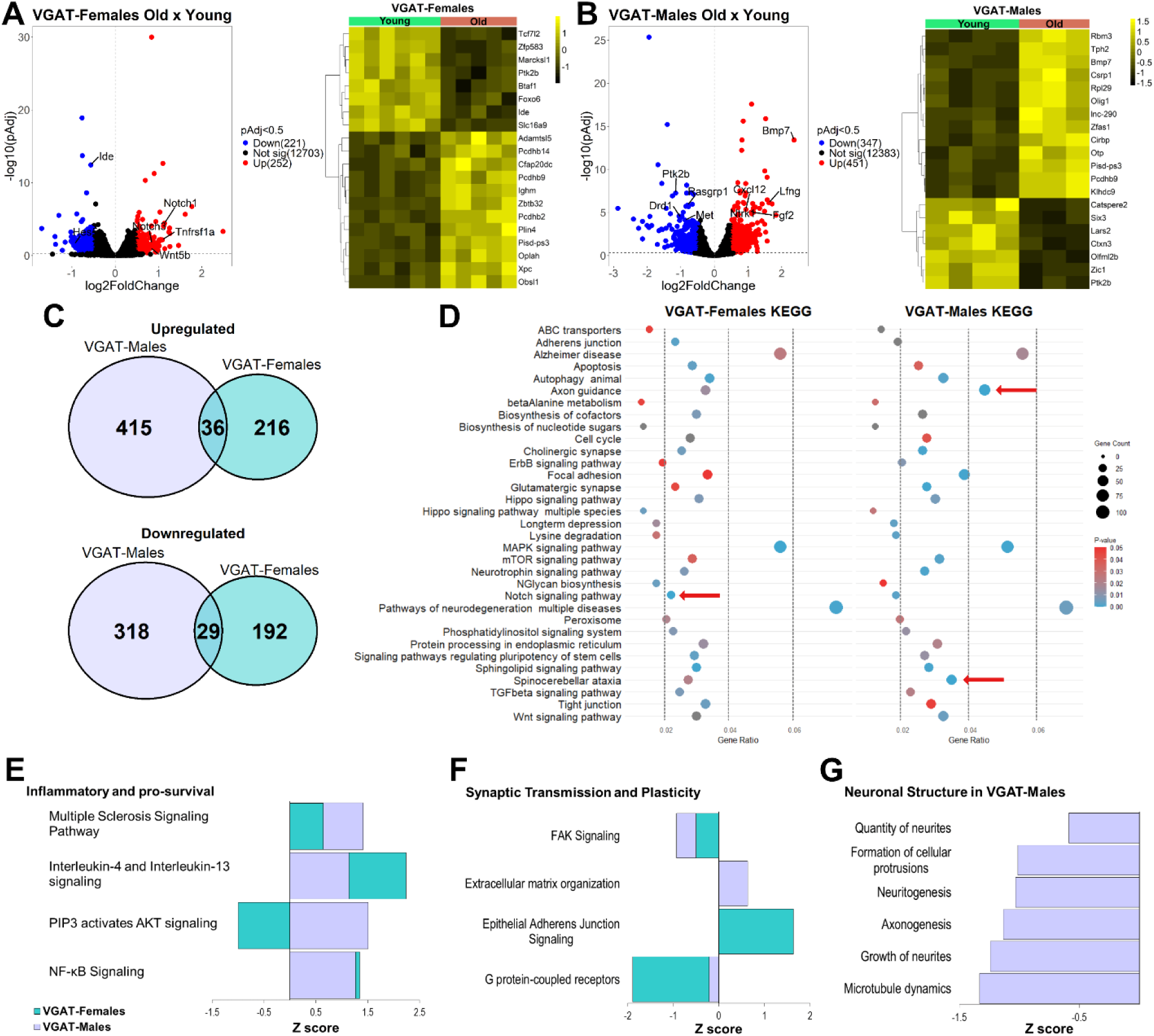
Sex-separate expression changes in GABA neurons with age. (A) Volcano plot of differential expression analysis for females and heatmap of the top differentially expressed genes (DEG; Cut off: Log2FC > |0.5|; Adjusted p value < 0.5). (B) Volcano plot of differential expression analysis for males and heatmap of the top DEGs. (C) Intersected up- and downregulated genes with age, between males and females. (D) Overall KEGG pathway analysis showing the most represented terms in both groups. Red arrows: Different changes in pathways. (E) IPA shows comparison of activated and deactivated paths for inflammatory and pro-survival signaling in both groups. (F) IPA analysis shows comparison of the downregulated pathways involved in synaptic transmission and plasticity for both groups. (G) IPA analysis shows predicted neuronal structure pathways downregulated for males, which were not altered in females. Pathways analysis cut off: p value <0.05 and q value <0.02.

As observed with DAT;NuTRAP mice, KEGG pathways analysis of the VGAT groups (Figure 6D) showed several overlapping pathways with similar gene ratios and p values. Also in agreement with DAT;NuTRAP data, the highest change in gene expression for both males and females was in genes connected to pathways of neurodegeneration (Figure 6D), corroborating what is known about aging being a factor in neurodegenerative diseases [126], but in a second type of midbrain neuron. Some pathways exhibited differences between the sexes; for example, males had considerably more altered genes related to spinocerebellar ataxia, similar to what was observed in dopamine neurons that had more ALS-related DEGs. Although there are no reports of higher male prevalence of spinocerebellar ataxia, spinocerebellar ataxia type 2 is clinically related to both PD and ALS [127–129], which are more prevalent in male patients. Other sex-biased pathways included axon guidance in males, and a higher gene ratio in females correlating with the Notch signaling pathway (Figure 6D), which is connected with AD (Kapoor and Nation, 2021) and has been shown to be activated in AD mouse models (Chen et al, 2018).

Pathway analysis showed that a few pro-survival pathways are activated with age in GABA neurons, for both males and females (Figure 6E). Curiously, one pathway that changed with age in opposite directions for males and females, the “PIP3 activates AKT signaling” pathway, is also an age-related sex-biased pathway in DAT;NuTRAP mice (Figure 5E). PIP3/AKT signaling has a variety of functions in neurons and is connected to both pro-survival signaling [79], and synaptic plasticity [130, 131]. Therefore, the exact effect of the activation or deactivation of PIP3/AKT in each cell type (dopamine or GABA neurons), or sex, with aging remains to be determined. IPA analysis also indicated that synaptic transmission and plasticity were affected in both males and females with age (Figure 6F), although these changes appear to be mostly similar between the two sexes. One last observation was that males presented changes in a variety of processes predicted to be involved in neuronal structures (Figure 6G), but these pathways were not significantly altered in VGAT-females (p value <0.05, pAdj <0.5).

These results suggest that while sexually divergent changes in gene expression occur with age in both dopamine and GABA neurons, they tend to converge into similar effects on biological processes. Overall, sex-effects appeared to be more prominent in dopamine than in GABA neurons.

## Discussion

While brain aging inevitably produces some decline in cognitive and behavioral function, whether it leads to the development of neurodegenerative disease [8–10] depends strongly on the individual. Further, while susceptibility to neurodegeneration and death can vary dramatically by cell type and brain region, the factors that are ultimately responsible for this variance are not well understood. Midbrain dopamine neurons and their regulatory circuits lie at the crux of some of the most common age-driven brain diseases [16, 21, 24, 25, 27]. GABA neurons in the same area may represent a more resilient neuron type, although inhibitory circuits do have suspected roles in brain disease [37, 38]. Here we sought a better understanding of the effects of normal aging on two distinct neuron types in a single brain area by investigating the transcriptomic changes in midbrain dopamine and GABA neurons. To do so, we used genetic crosses to create and validate two neuron type-specific NuTRAP lines (DAT;NuTRAP and VGAT;NuTRAP). The presence of GFP ribo-tags expressed only in cells containing *Cre recombinase* allowed for the acquisition of neuron type-enriched RNA-Seq datasets (Figure 1). Of note, the NuTRAP construct also supported mCherry expression on the nuclei of Cre-positive neurons, which was not the subject of the current study but will allow for epigenomic studies in the future. Overall, the present findings illuminate some of the common and divergent pathways of age-driven molecular alterations in both neuronal types, offering new insights into the mechanisms underlying neuronal vulnerability and resilience in neurodegenerative contexts.

Dopamine neurons exhibited extensive transcriptional changes with aging (Figures 3 and S1), including the upregulation of response to inflammation and pro-survival pathways (e.g., MAPK, PI3K/Akt and Notch signaling) and the downregulation of synaptic and cell signaling pathways (e.g., calcium, cAMP, apelin signaling). The combination of up- and downregulation events reflect a global change in neuronal homeostasis, an effect common to aged neurons observed in other regions of the brain. Although several of the findings observed here are consistent with those reported elsewhere [44, 76, 78, 113], their presence in dopamine neurons likely has broad consequences due to their regulatory role in a variety of functional processes. Moreover, we here and others [4] observed aging alterations in mitochondrial gene expression, a known outcome [132] that can prompt dopaminergic dysfunction and is often detected in patients and animal models of PD [87, 89, 91]. Overall, the molecular events involved in dopamine neuron aging provide an important link between age-related dysfunction and the development of neurodegenerative diseases such as PD and AD [41, 92, 109, 133]. Notably, the downregulation of synaptic genes aligns with literature showing age-related epigenetic changes as key factors affecting synaptic structure and function during aging [134].

When compared to dopamine neurons, GABAergic neurons showed fewer transcriptional changes, which may reflect to some extent their historically hypothesized resilience (Rissman et al., 2007). One noteworthy alteration observed with aging in GABA neurons is the upregulation of serotonergic signaling pathways and synapses, which is consistent with reports in other brain regions [45, 101]. In particular, the increased expression of the gene *Cyp2d22*, which codifies one of the enzymes involved in the synthesis of serotonin, has been reported to be involved in an alternative serotonin synthesis pathway [135]. Although there is a consensus of the overall importance of the effects of serotonergic signaling in the aged brain, particularly in AD patients [136], the individual events that occur are more controversial [137], likely due to the different responses observed across areas of the brain [138]. Overall, it appears that with aging there is a global decrease in 5-HT receptors and an increase in serotonin turnover [137], which is consistent with the increased synthesis of serotonin-related enzymes observed in GABA neurons (Figure 3H and S1E). On the other hand, the increased expression of these enzymes (Tph2, Pla2g4e and Cyp2d22) also correlates with the increase of serotonergic signaling in the presence of inflammation and cellular stress [102–104], another event we observed in aged GABA neurons (Figures 3F and 3G). Whatever the driving factors, serotonergic alterations certainly influence the excitatory and inhibitory inputs regulated by GABA neurons and contribute to age-related cognitive decline [39].

Concerning the similarities in the aging of GABA and dopamine neurons, both neuronal types demonstrated overlapping pathways associated with increased response to inflammation, and declining of synaptic structure and plasticity. As chronic inflammation and cellular senescence have been identified as two hallmarks of aging [139], it might be expected that aged dopamine and GABA neurons would also express genes connected with cellular senescence. Changes in Notch, MAPK, cAMP and cytokine signaling were observed in both cells (Figures 3 and 4) and are signatures of the senescence-associated secretory phenotype [83]. In association with changes in calcium homeostasis and synaptic structure and plasticity, these alterations point towards the occurrence of neuronal “senescence” in both (postmitotic) cell types [83, 126]. These shared changes highlight some of the potential mechanistic links between neuronal aging and the increased vulnerability of these cells, particularly dopamine neurons, to neurodegeneration. Further, some common findings could reveal potential targets for prevention therapy that could modulate inflammation and/or reinforce synaptic plasticity pathways [140–142].

Despite the undeniable evidence connecting age-related gene expression changes to deficits in neuronal function and neurodegeneration, our data also highlighted several mechanisms of adaptation that allowed dopaminergic and GABA neurons to maintain homeostasis during healthy aging. In fact, to maintain proper function, neurons in the aging brain knowingly undergo several changes in metabolism and connectivity [143]. One such change is likely the increase in PI3K/Akt signaling pathway we observed in DAT;NuTRAP mice (Figure 3C), which activates pro-survival signaling cascades during stress [79, 80]. Mitochondrial function adaptation is another mechanism previously observed in aged neurons [88] that we identified here in dopamine neurons, particularly in aged males that showed upregulation of genes involved in complex IV (Figure 5H). Dopaminergic neurons in females showed increased upregulation of genes involved in protection against oxidative stress (Figure 5H) [122] and are likely an adaptive mechanism that confer defense of these cells against mitochondrial dysfunction. Furthermore, changes in synaptic connectivity and plasticity could also represent adaptive mechanisms during healthy aging, since observations suggest that reduced synaptic gene expression predicts longer lifespan among healthy individuals [143, 144]. Some of the changes in calcium dynamics and signaling (Figures 3C, 3K-L and Figures 5H-J) are likely compensatory mechanisms promoted by these neurons as an attempt to decrease the levels of circulating intracellular calcium, particularly in situations of mitochondrial dysfunction [44, 145].

Concerning sex differences in the age-related transcripts, we found that these were prominent in autosomally-encoded genes, and were observed in both dopamine and GABA neurons. As age-related changes in estrogen and estrogen receptor levels promote alterations in female brains that affect synaptic connectivity, neuronal function, and gene expression [146, 147], the detection of sex differences was anticipated. In fact, sex differences in gene expression are often conserved across species [148] and several areas of the brain [149, 150], particularly during aging. Although dopamine neurons of the SNc appear to have fewer sex differences in gene expression patterns than other regions of the brain [48], age-driven sex effects in dopaminergic function and physiology have been reported [20, 114, 151]. Some of these events were reflected as altered gene expression particularly in males, such as decreased glutamate receptor signaling, potassium channel expression (Figure 5E), alterations in mitochondrial proteomics and bioenergetics (Figure 5H) and alteration in calcium dynamics (Figure 5H-I). Interestingly, both dopamine and GABA datasets presented sex differences that appeared to match up with the known prevalence of neurodegenerative diseases [46–48]. Alterations in dopamine neurons connected to PD and ALS were also reflected in the increased of gene expression changes connected with spinocerebellar ataxia in GABA neurons (Figures 5D and 6D), which could reflect similarities in aging and susceptibility to neurodegeneration in the two cell types. Furthermore, some of the female findings such as altered Notch signaling of GABA neurons (Figure 6E) could reflect the connection of the pathway to AD prevalence in females [48, 77, 78]. Curiously, although females undergo major brain-related hormonal and systemic changes with age [146], the present work (Figures 5G and 6G) and published data concerning sex effects on neuronal aging [20, 114] suggest that midbrain dopamine and GABA neurons in males are more sensitive to age-driven changes than females. As the expression of tyrosine hydroxylase (the rate-limiting enzyme for production of dopamine) is dependent on the presence of Sry protein in a subset of male dopaminergic neurons [152], it is possible that some of the age-driven sex differences observed in the midbrain of males could be connected to loss of Sry expression with aging. Furthermore, studies of the hippocampus and a few other regions detected that male brains have globally decreased anabolic and catabolic capacity, and are therefore more susceptible to neurodegeneration [150]. Hence, the sex differences we observed between dopamine and GABA neurons with aging warrant further investigation, and should be considered when studying the effects of aging on the midbrain and related circuits.

One caveat of the current work is that the quasi-bulk nature of the NuTRAP approach did not allow for a further understanding of specific dopaminergic subpopulations [153]. Future studies could integrate spatial transcriptomics in the midbrain or single-nuclei analyses to further unravel the complexity of age-driven changes in dopamine and GABA neurons, as has been observed in other regions of the brain [4, 14, 15]. Further, Patch-sequencing of dopamine neurons was recently demonstrated in an AD model [27], and a similar approach could be used here to study subpopulations of physiologically-characterized dopamine neurons with aging. Functional studies will also be necessary to validate the roles of identified pathways in neuronal aging and disease progression.

In summary, the present work reveals molecular underpinnings of neuronal aging, highlighting shared and distinct changes in dopamine and GABA neurons while providing insights into mechanisms that could link aging to neurodegeneration. These findings offer a framework for understanding how aging shapes dopamine and GABA neuron function and susceptibility to neurodegenerative diseases, emphasizing the potential of targeting age-associated pathways for therapeutic interventions.

## Acknowledgements

The authors would like to acknowledge the OMRF Clinical Genomics Center for sequencing services and assistance with experimental planning and strategies. The authors also acknowledge the OMRF Center for Biomedical Data Sciences for training and brainstorming sessions involving data analysis, as well as creation and maintenance of the RNA-Seq pipeline used for fastq files pre-alignment processing, genome alignment and post-alignment counts, and the DESeq2 (R) pipeline used for differential analysis. We also acknowledge Stuart Glenn for his help with file accessibility and maintenance of the OMRF High Performance Computing (HPC) cluster.

## Author Contributions

ALDB designed and performed the research, processed TRAP samples, prepared libraries, analyzed the data and wrote the manuscript. HEB designed and performed the research, processed TRAP samples, prepared libraries, analyzed the data and revised the manuscript. KDP helped with TRAP processing and library preparation. KAC performed animal husbandry and genotyping for all animals used in the study. WMF designed the research, revised and edited the manuscript. MJB designed and performed the research, supervised experiments and analysis, revised and edited the manuscript, and is responsible for the primary funding used for this study.

## Supplemental material

**Table S1 –** Primers used for used for mouse genotyping and antibodies used for immunofluorescence.

**Table S2 –** Lists of genes used to verify dopaminergic or GABAergic enrichment.

**Table S3 –** Orthogonal analysis compared with Hahn et al. (2023) [14].

**Table S4 –** Orthogonal analysis compared with Kilfeather et al. (2024) [4].

**Table S5 –** Differential expression results for all groups. DAT;NuTRAP (grouped and sex-separate) and VGAT;NuTRAP (grouped and sex-separate).

